# The duration of chronic restraint stress protocols does not predict effect sizes in behavioural tests: a meta-analysis

**DOI:** 10.64898/2026.03.27.714712

**Authors:** Nicola Romanò, John Menzies

**Affiliations:** Institute of Neuroscience and Cardiovascular Research, University of Edinburgh, Edinburgh, UK; Edinburgh Medical School, University of Edinburgh, Edinburgh, UK; University of Edinburgh-Zhejiang University Joint Institute, Zhejiang University School of Medicine, Zhejiang University, Haining, China

## Abstract

Stressors are commonly used to induce models of anxiety or depression in rats. The effectiveness of these stressors is often evaluated using behavioural tests. In a previous meta-analysis of chronic variable stress (CVS) procedures, we hypothesised that the duration or intensity of CVS procedures would correlate with the magnitude of the effect sizes reported in behavioural tests. However, we did not find strong evidence for this correlation, and were concerned that variability in CVS procedure design, both the duration and the types of stressors used, may have made it challenging to determine which factors influenced effect size. To address this, we carried out a meta-analysis of studies using a simpler stress procedure – chronic restraint stress (CRS) – to study the relationship between the duration of CRS procedures and the effect sizes obtained in subsequent behavioural tests. In articles using CRS procedures with rats, we documented the duration of restraint and carried out a meta-analysis of four behavioural tests: the forced swim test (FST), the sucrose preference test (SPT), the elevated plus maze (EPM) and the open field test (OFT). We found that chronic restraint stress increased immobility in the FST, decreased sucrose preference in the SPT, decreased time spent in the open arms of the EPM but had no effect on time spent in the centre of the OFT. However, the effect sizes in all behavioural tests were not moderated by the duration of the CRS procedure, indicating that longer CRS procedures are not associated with larger effect sizes.

## Introduction

Studies that impose stressors on laboratory animals are widely used to induce psychological and/or physiological states that are believed to represent models of human anxiety and depression. However, there is controversy around whether some translational approaches are, in fact, valid models of human disease, and whether these models have predictive value (Worp *et al*., 2010; Martić-Kehl *et al*., 2012; Pound & Bracken, 2014; McGonigle & Ruggeri, 2014; Pound & Ritskes-Hoitinga, 2018; Ferreira *et al*., 2020; Stanford, 2020; Gencturk & Unal, 2024). Several families of stress procedures have been developed to model different aspects of human stress, including chronic variable stress (CVS), social defeat stress and pre-and peri-natal stress (Petković & Chaudhury, 2022). CVS involves relatively complicated procedures comprising repeated and multiple exposures to stressors including cold or heat, the removal of food, water or bedding, exposure to hypoxia, noise, stroboscopic lighting or predator odours, social isolation or overcrowding, tilting or shaking the home cage, forced swimming, or painful stimuli (Willner, 2017; Antoniuk *et al*., 2019). Accordingly, CVS procedures are often perceived as time-consuming, resource-intensive and having a significant negative impact on the welfare of animals and on the well-being of people working with and caring for the animals (King & Zohny, 2022; Morahan *et al*., 2024). In contrast, CRS normally involves a single type of stressor: placing rodents into a confined space that restricts their ability to move, doing so without deliberately imposing pain or discomfort, normally imposed daily for consecutive days. It is commonly believed that CRS results in physiological and behavioural changes that represent a model of anxiety and/or depression (Petković & Chaudhury, 2022), though some studies show decreased anxiety-and depression-like behaviour in after CRS procedures, albeit with very short (5 min) bouts of restraint in mice or rats (Parihar *et al*., 2011; Lee *et al*., 2021), and repeated exposure to the same restraint conditions can lead to habituation, with animals exhibiting attenuated physiological or behavioural stress responses over time (Buynitsky & Mostofsky, 2009).

Behavioural change induced by stress is often evaluated using a panel of behavioural tests: the forced swim test (FST; where an increased time spent immobile is deemed to reflect a depressive-like state), the sucrose preference test (SPT; where a decreased preference for sucrose over water is deemed to reflect an anhedonic state), the elevated plus maze (EPM) and the open field test (OFT; where, in both tests, a decreased time spent in open, exposed areas is deemed to reflect increased anxiety). There are questions about the reliability and validity of these tests (Kara *et al*., 2018; Stukalin *et al*., 2020; Molendijk & Kloet, 2022; Rosso *et al*., 2022; Primo *et al*., 2023; Berrio *et al*., 2024), but they are often used as evidence for stress-induced behavioural changes and for the effectiveness of interventions that alleviate anxiety-and depression-like behaviours.

In a previous study, we systematically explored whether differences in the duration, intensity and burden of a range of rat CVS procedures were associated with differences in effect sizes in the FST, SPT, EPM and OFT (Romanò & Menzies, 2025). In that study, we reasoned that if a change in behaviour reflects a biological and/or psychological consequence of CVS, then longer or more severe CVS procedures may be associated with larger effect sizes in behavioural tests. However, we found only weak correlations between the intensity of stressors applied and the effect sizes obtained in subsequent behavioural tests. Here, we set out to explore the same question in CRS studies; a stress procedure that is, on the face of it at least, far simpler and less variable than CVS procedures, thus a more amenable context to study the effect of the length of exposure to stress on behavioural outcomes. CRS has been likened to “a continuous, predictable stress people experience every day” such as the “daily repetition of a stressful job, social or financial stress, familial stresses, or day to day stresses that are repeated and constantly stacked upon the previous day’s workload” (Wang *et al*., 2017). The implication being that each exposure to stress may be additive in terms of the overall negative effect on the animal over time. This intuitive interpretation of CRS as an increasing allostatic load seems plausible to us, so we hypothesised that longer CRS procedures would be associated with a larger magnitude effect sizes in post-CRS behavioural tests. To explore this, we extracted data from articles reporting behavioural testing after a CRS procedure using rats and performed a meta-analysis to explore whether the length of the CRS procedure influenced the effect sizes obtained in behavioural tests.

## Materials and methods

We did not pre-register this study. MEDLINE, PubMed Central, Web of Science and Embase were searched using the term ‘“chronic restraint stress” AND “rat”’ in all fields. The search was done on 7 June 2026 and included all articles published from 2015 to 2025. Exclusion criteria are shown in Figure 1.

**Figure 1:**
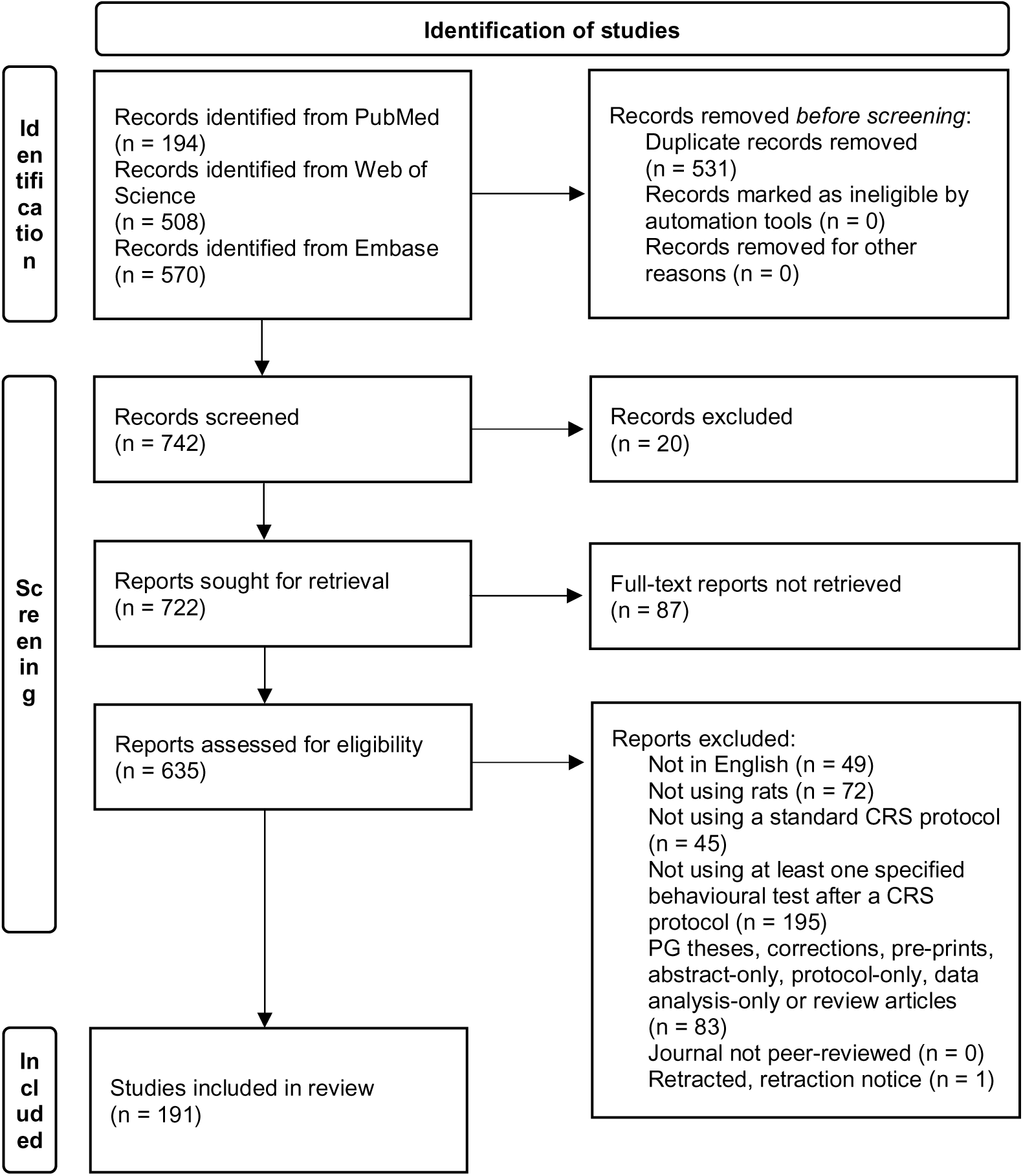
Identification of studies (following Page et al., 2021). The search term was ‘“chronic restraint stress” AND “rat”’. MEDLINE, PubMed Central, Web of Science and Embase were searched on 7 June 2026. Thirteen records were excluded during screening because they comprised links to an individual article figure, and seven records were excluded because they were published outside of our study’s time range.

For each article, we used the information provided in the Methods sections to document the duration of the restraint session, the number of restraint sessions used each day, and the total duration of the CRS procedure in days. We also documented the strain, sex and age and/or body weight of the rats used, and the timing of the restraint session with respect to the light-dark cycle (these data are available at https://github.com/nicolaromano/Romano_Menzies_2026_CRS). We did not use these metadata in our analysis, because age/weight were normally given in what appeared to be an estimated range rather than a specific range of values, only 9% of the included studies used females, and information on the timing of the restraint session was often not given or ambiguous.

We documented the characteristics of four types of behavioural tests used to evaluate the effects of CRS: the FST, the SPT, the EPM and the OFT. For the FST, we documented the duration of the test. For the SPT, we documented the concentration (%) of sucrose used, the duration of exposure to the sucrose/water choice and whether and for how long the animals were deprived of food and/or water before the test. For the OFT, we documented the duration of the test and the time spent in the central part of the field. For the EPM, we documented the duration of the test and the time spent in the open arms.

To extract the reported outcomes from data figures, we used an online tool (foxyutils.com/measure-pdf) to measure the size of the mean and its error bar (representing either the standard error of the mean (SEM) or the standard deviation (SD)) in relevant figures. If the article gave the SEM, we calculated the SD using the reported sample size. If the sample size was given as a range, we used the lowest number in the range to calculate the SD. Some articles reported % sucrose preference in a bar graph with the % preference axis labelled from 0 to 1. In those instances, we assumed the axis had been mislabelled and the data reported ran from 0% to 100%.

Meta-analysis was conducted using the metafor package (version 4.8.0; Viechtbauer, 2010) in R 4.3. Effect sizes were calculated using small sample size-adjusted standardized mean differences (Hedges’ g) using the following calculation:

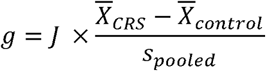

Where the small-sample correction factor J (Hedges, 1981) is defined as:

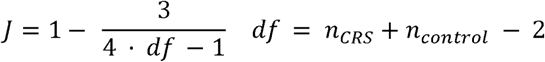

And the pooled standard deviation s_pooled_ is defined as:

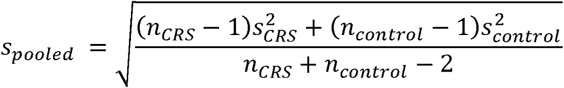

These calculations result in a positive effect size in the FST when immobility increases; a negative effect size in the SPT when sucrose preference decreases, in the EPM and OFT when time in the open arms or in the centre decrease respectively. A random-effects multilevel meta-regression model was fitted using restricted maximum likelihood for variance estimation; the Knapp-Hartung adjustment for standard errors and p-value was used to account for small sample sizes. The article’s randomly-allocated ID number was used as a random effect to control for studies that reported multiple outcomes (for example, when data from multiple CRS durations were reported in a single article). The total duration of CRS and, in the case of the SPT, the duration of prior water and food deprivation was used as a moderator on the effect sizes. For the SPT, all studies deprived animals of water and food at the same time and for the same duration, aside from three studies where only water was removed (Meng *et al*., 2021; Arab *et al*., 2023; Elfakharany *et al*., 2024), but we did not account for this difference in our analysis. Heterogeneity was quantified using the I^2^ statistic (Higgins & Thompson, 2002). Small study effects were evaluated through visual inspection of funnel plot asymmetry. Funnel plots were generated using

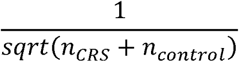

as study precision, to reduce distortion associated with standardized mean differences (Zwetsloot et al., 2017). As in our previous study (Romanò & Menzies, 2025), we calculated a pairwise Pearson correlation between for all possible pairs of observations to ask whether effect sizes correlated between different behavioural tests.

To determine whether any specific article had a disproportionate influence on the estimated effects, we conducted a leave-one-out sensitivity analysis (Viechtbauer & Cheung, 2010), by sequentially refitting the model after removing each study in turn and comparing the resulting predicted effects to the full model.

No generative AI tools were used in the data extraction or analysis, or in the preparation of the manuscript.

## Results

We systematically documented and analysed a total of 216 experiments from 191 articles where at least one of the FST, the SPT, the EPM and/or the OFT were performed after imposing CRS in rats. Detail of the rats used, and CRS protocol design are available at https://github.com/nicolaromano/Romano_Menzies_2026_CRS. One hundred articles (52%; all percentages are given as the percentage of the total number of articles meeting the inclusion criteria) used Wistar rats, including Wistar Han, Wistar-Crl:WI and Wistar Kyoto strains. Eighty-eight articles (46%) used Sprague-Dawley rats, two articles (1%) used Long-Evans rats. The “Spontaneously Hypertensive Rat”, the Flinders Sensitive Line and the TgF344-AD strain were each used in one article. One article used both F334 and Wistar Kyoto rats. One article did not report the strain used.

One hundred and seventy-four articles (91%) used males only, seven articles (4%) used females only, with two of those articles using pregnant females. Ten articles (5%) used both male and female rats. Three articles did not report the sex of the rats used. One hundred and eighty-three articles (96%) reported the age and/or initial body weight of the rats. All studies that reported using adult rats (median age: 8 weeks, range 4-52 weeks; median initial body weight: 210 g, range 80-415 g).

All but one article reported detail of the CRS protocol used. All but three articles (98%) reported using only one restraint session per day, and that the restraint session was of a fixed duration. There was variability between articles in the duration of each daily restraint session (median: 6 h; range: 20 min-8 h), and in the duration of the entire procedure (median: 21 days; range: 1-63 days; Figure 2). One article reported using three 45-min sessions daily, one article reported using two 1-hour sessions daily, and one article reported alternating between two 1-hour sessions and one 2-hour session daily in an effort to limit rats’ adaptation to the stressor. One article reported using one daily 1-hour session for 14 days, then escalated to one daily 6-hour session for an additional 14 days. An explanation for the escalation was not given in the article. Most articles reported imposing restraint on consecutive days, but three articles reported imposing restraint on Mondays to Fridays only, and one article reported imposing restraint every other day. We calculated the total duration of restraint by multiplying the duration of the daily session by the total number of days on which restraint was imposed. The median total duration of restraint was 84 hours (range: 2-378 hours).

**Figure 2:**
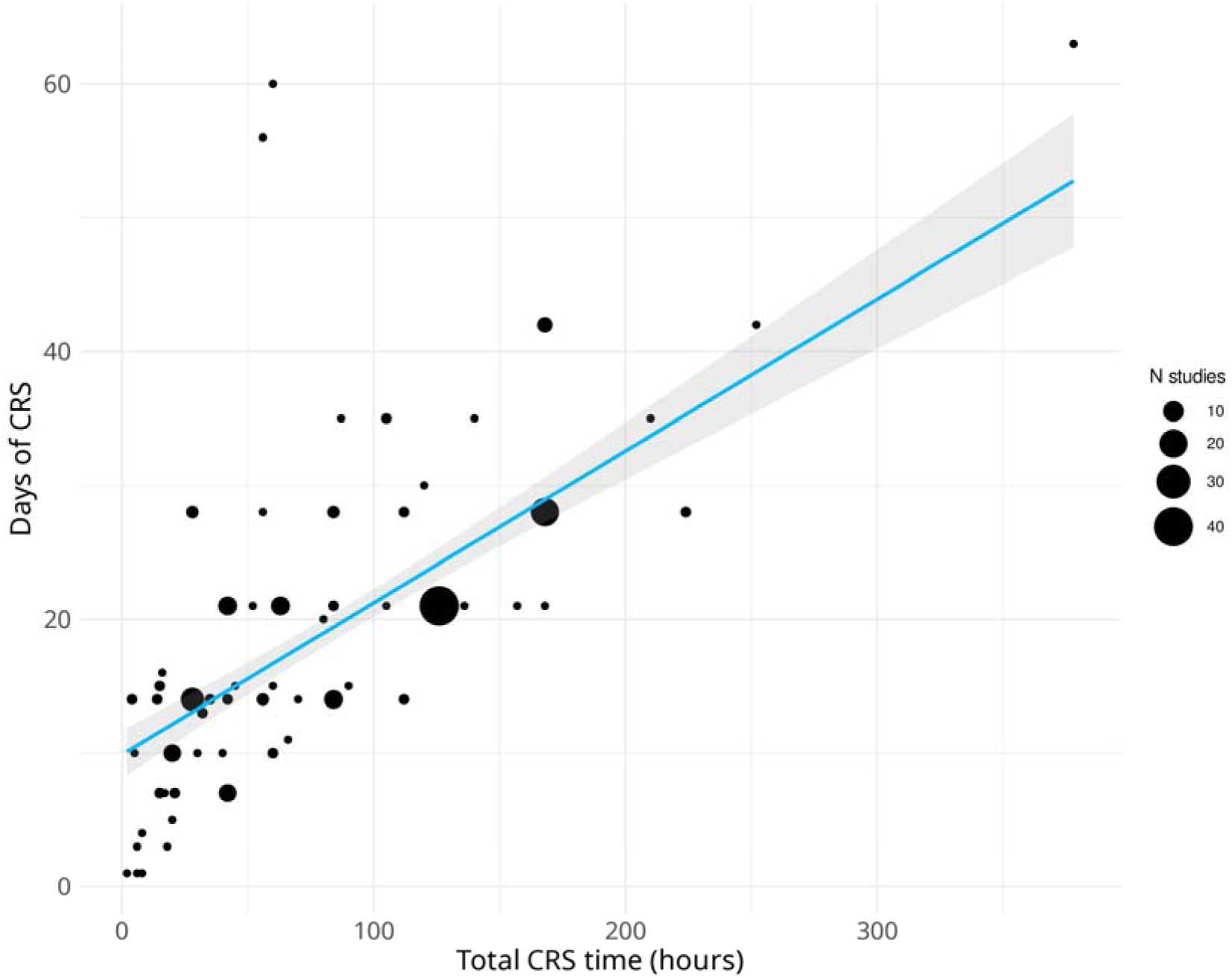
Scatter plot of the total time restrained (hours) versus duration of the entire procedure (days). The size of each point is proportional to the number of studies that used the specific parameters. The blue line is the linear regression (β_total time_=0.123, t_65_=11.29, p<0.001, adjusted R^2^=0.66) and the grey band show 95% confidence intervals.

A range of restraint devices was used. Most articles (123 articles; 64%) used a plastic tube, bottle, cage or box. Thirty-one articles (16%) used a wire mesh, thirteen articles (7%) used a flexible conical plastic bag, and six articles (3%) used “soft bands”. Three articles used a metal cylinder, two used a “barbed wire mesh”, one used a wooden box, one used an “elastic immobilizer”, one used a “custom-designed press plate” and one used a “glass restrainer”. Forty-two articles (22%) did not report detail of the restraint device used. One article reported two studies using different devices: a wire mesh to “restrain” or a plastic bag to “immobilise” the rats (Ampuero *et al*., 2015).

Ten articles (5%) reported carrying out behavioural testing after more than one CRS procedure (of differing durations) or at more than one point during a CRS procedure. One study reported using saccharin rather than sucrose in the SPT.

Because rats display circadian activity (Cuesta *et al*., 2009), and there are circadian variations in rats’ stress axis activity (Allen-Rowlands *et al*., 1980; Atkinson & Waddell, 1997), we decided it would be valuable to document when during the light-dark cycle the restraint sessions took place. However, only fifty-four articles (28%) reported sufficient information to determine unambiguously whether the restraint sessions took place in the light or dark phase. Eight articles (4%) carried out the restraint sessions during the dark phase and forty-six articles (24%) carried out the restraint sessions during the light phase.

For each behavioural test, the range of ‘total duration of restraint’ values are given in Table 1 alongside the total number of included articles, the total number of experiments in those articles and the total number of excluded experiments. Experiments were excluded because they either did not report the number of rats, did not provide a the mean and/or a measure of variability, did not report the length of CRS, or did not report the length of water deprivation before carrying out an SPT.

**Table 1:** Number of experiments where effect size estimates extracted from articles using each type of behavioural test, and the range of total duration of restraint used (hours) prior to testing.

| Test | Total number of articles | Total number of experiments | Number of excluded experiments | Total duration of restraint (range, hours) |
| --- | --- | --- | --- | --- |
| FST | 93 | 106 | 30 | 1.6-336 |
| SPT | 73 | 107 | 55 | 15.8-378 |
| EPM | 77 | 91 | 28 | 5-180 |
| OFT | 116 | 146 | 120 | 4.7-180 |

We noted an unexpected breadth in the types of behavioural measurements reported in the OFT. A total of thirty-two different measurements were reported across 109 articles, with a median of three different measurements reported per article (range: 1-7; these data are available at https://github.com/nicolaromano/Romano_Menzies_2026_CRS). The OFT is normally used to evaluate locomotion and anxiety, with a lower amount of time spent in the centre of the OFT believed to reflect a higher level of anxiety (Kraeuter *et al*., 2019; Võikar & Stanford, 2023). However, only thirty-two articles using the OFT (29%) reported the time spent in the centre. The most frequently reported measurements were the total distance travelled (reported in 58% of articles), the number of lines/squares/grids crossed (41%) and the number of times the rat reared (43%).

We conducted a meta-analysis of the estimated effect sizes in each of the four behavioural tests. Across 67 independent studies (76 effect sizes), CRS was associated with a significant increase in immobility in the FST (Hedges’ g = 2.12, 95% CI [1.50, 2.74], t_74_=6.809, p=2.23 x 10^-9^, calculated at the median value of CRS exposure: 105 hours). The total duration of CRS had a non-significant moderating effect on immobility (β_length_[95% CI] = 0.0010 [-0.0055, 0.0075], t_74_ = 0.3011, p = 0.7642) (Figure 3). Between-study heterogeneity was very high (I^2^ = 86.9%). Visual inspection of the funnel plot (Figure 4) suggested some asymmetry, with greater dispersion and a tendency towards positive residuals among smaller studies; the asymmetry appeared to be driven in part by one extreme positive residual. (Zhu *et al*., 2021).

**Figure 3:**
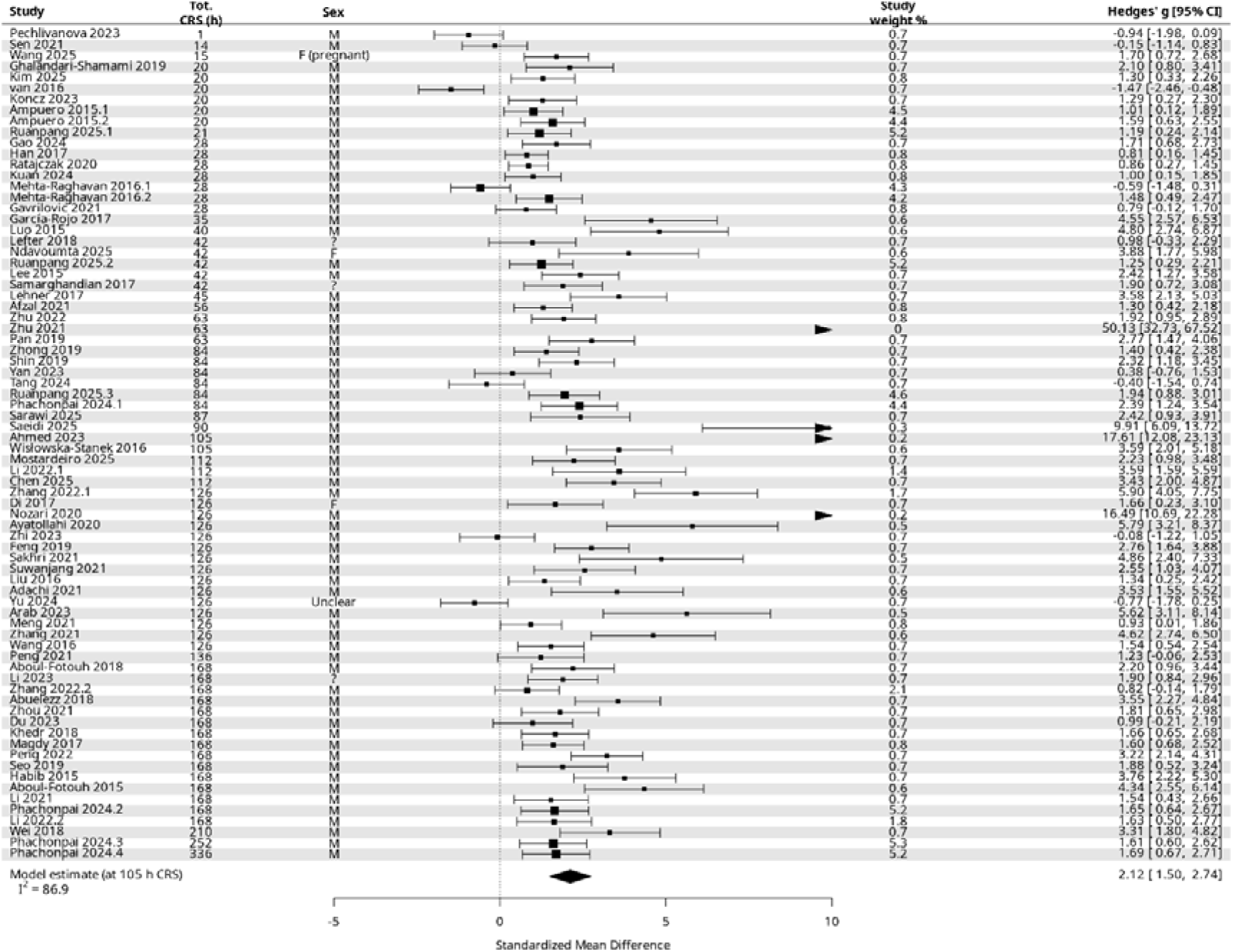
Forest plot of effect sizes extracted from studies measuring immobility time using an FST.

**Figure 4:**
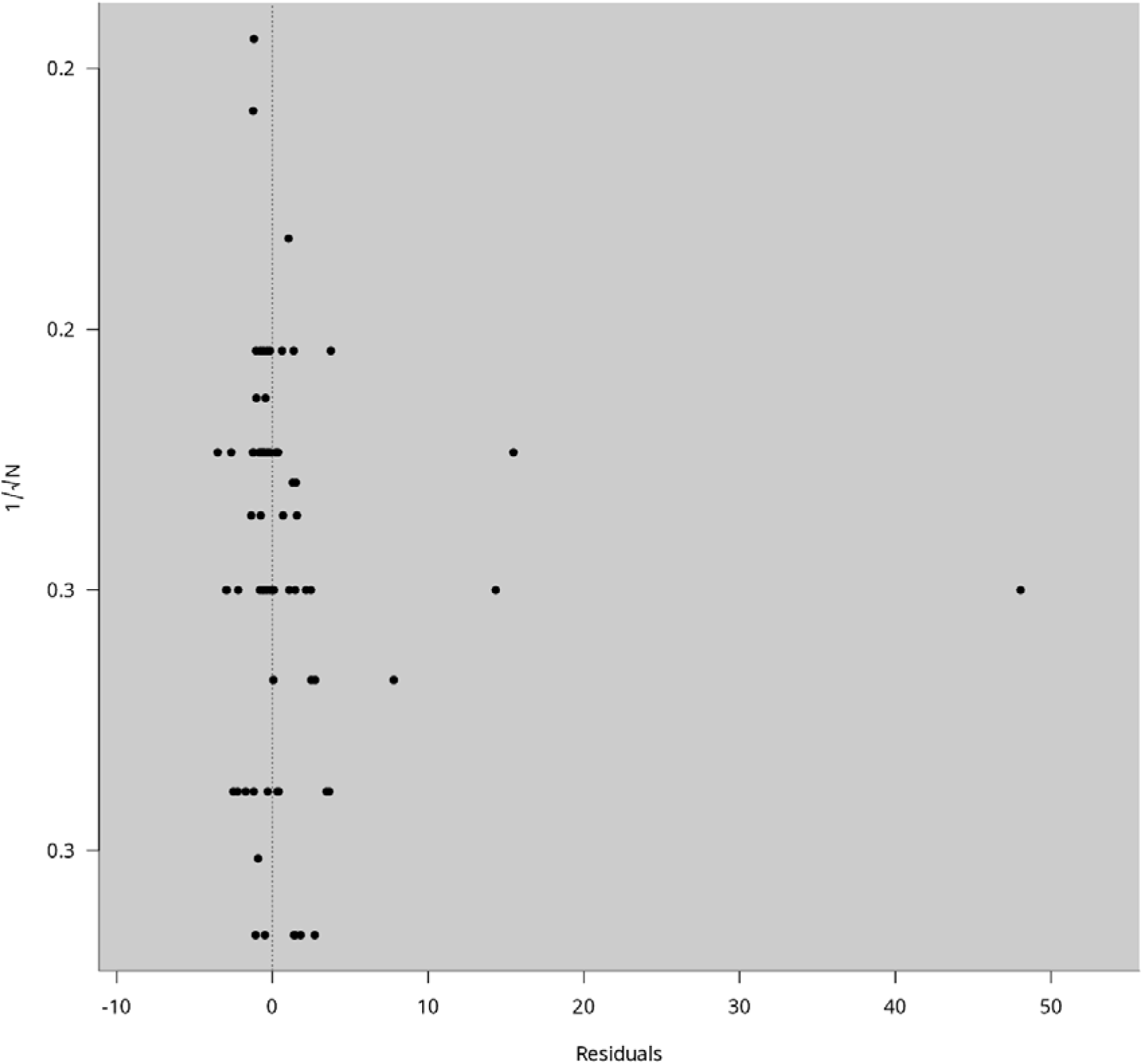
Funnel plot depicting effect size and study precision from studies measuring immobility time using an FST.

Across 31 independent studies (52 effect sizes), CRS was associated with a decrease in sucrose preference in the SPT. Predictions at the median CRS length (126 hours) indicated a reduction in sucrose preference both with no water or food deprivation (Hedges’ g =-1.704, 95% CI [-2.80,-0.61], t_49_=-3.136, p=0.003) and with 24 hours water and food deprivation (Hedges’ g =-1.611, 95% CI [-2.26,-0.96], t_49_=-5.009, p=7.5 x 10^-6^). CRS duration did not significantly moderate effect size (β_length_[95% CI] =-0.0032, t_49_ =-1.35, p = 0.185). Water and food deprivation also did not significantly moderate the effect (β_water_[95% CI]-0.0039 [-0.0528, 0.0606], t_49_ =-0.137, p = 0.89) (Figure 5). Between-study heterogeneity was high (I^2^ = 76.2%). Visual inspection of the funnel plot (Figure 6) showed little evidence of a small study effect.

**Figure 5:**
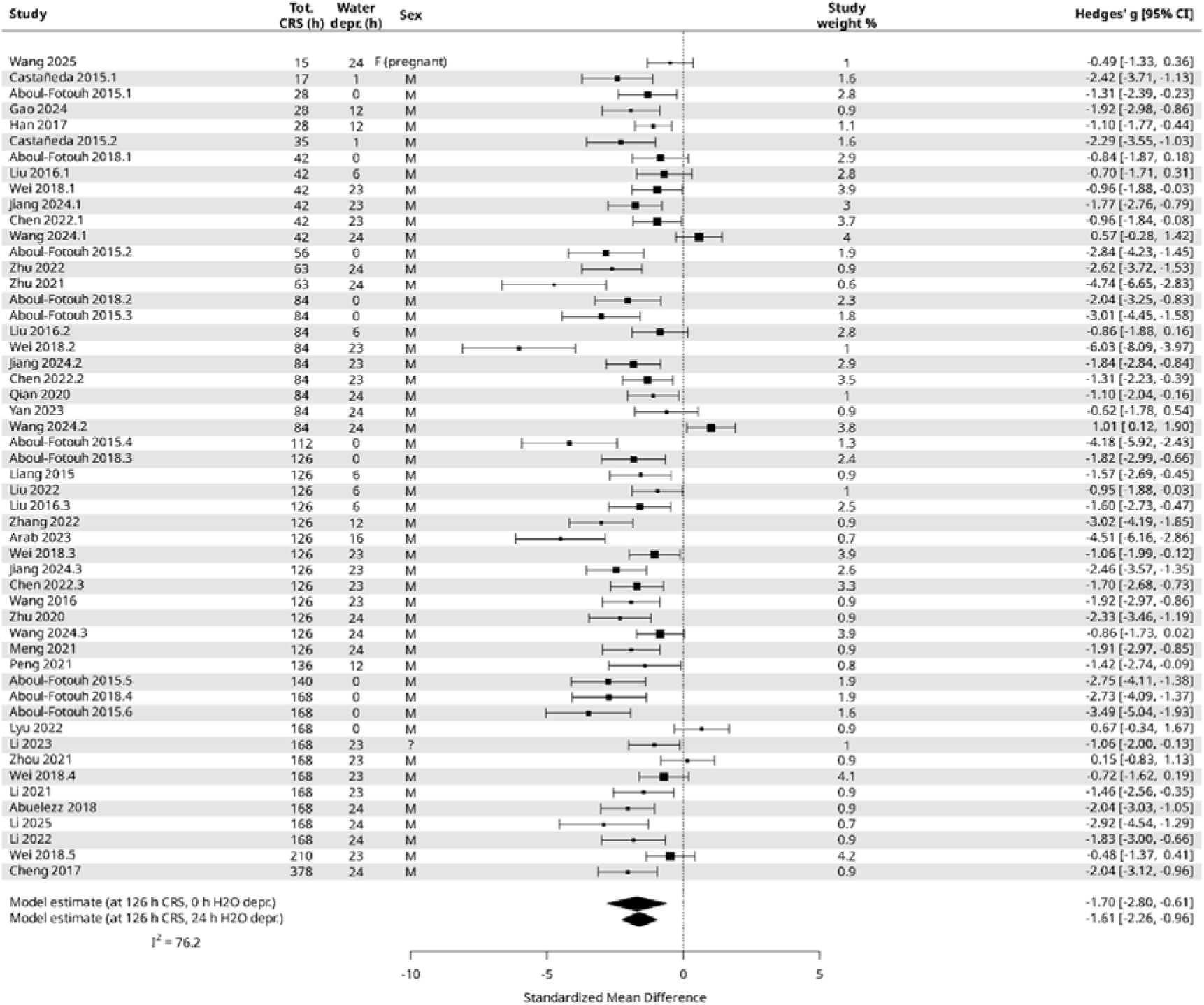
Forest plot of effect sizes extracted from studies measuring sucrose preference using an SPT.

**Figure 6:**
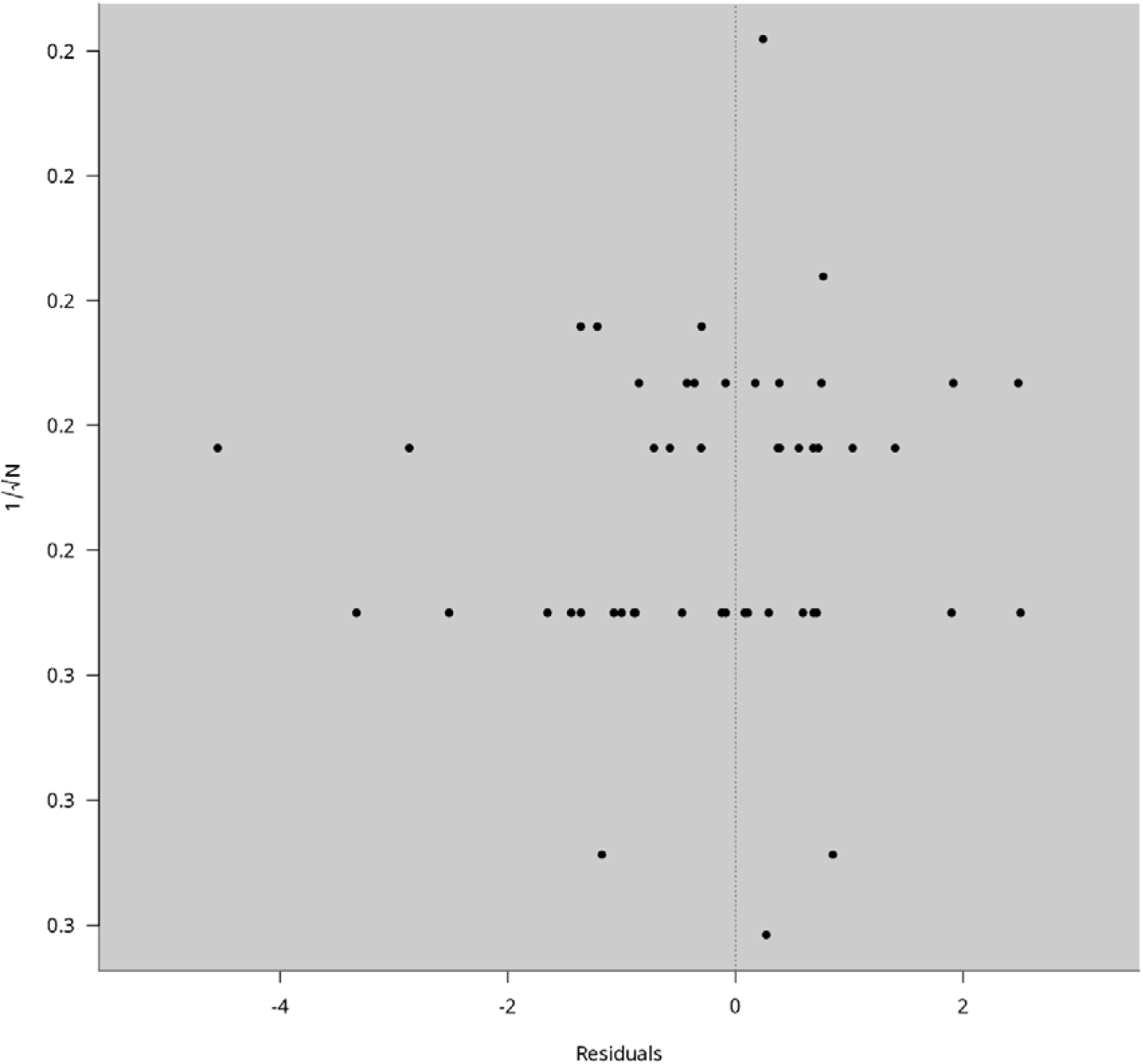
Funnel plot depicting effect size and study precision from studies measuring sucrose preference using an SPT.

Across 22 studies (26 effect sizes), CRS was associated with a non-significant decrease in time spent in the centre of the OFT (Hedges’ g =-1.104, 95% CI [-2.42, 0.22], t_24_=-1.726, p=0.10, calculated at the median CRS length: 87 hours). The total duration of CRS had a non-significant moderating effect on time spent in the centre (β_length_[95% CI] =-0.0106 [-0.0228, 0.0016], t_24_ =-1.789, p = 0.086) (Figure 7). Between-study heterogeneity was very high (I^2^ = 96%). Visual inspection of the funnel plot (Figure 8) did not show clear evidence of small-study effects.

**Figure 7:**
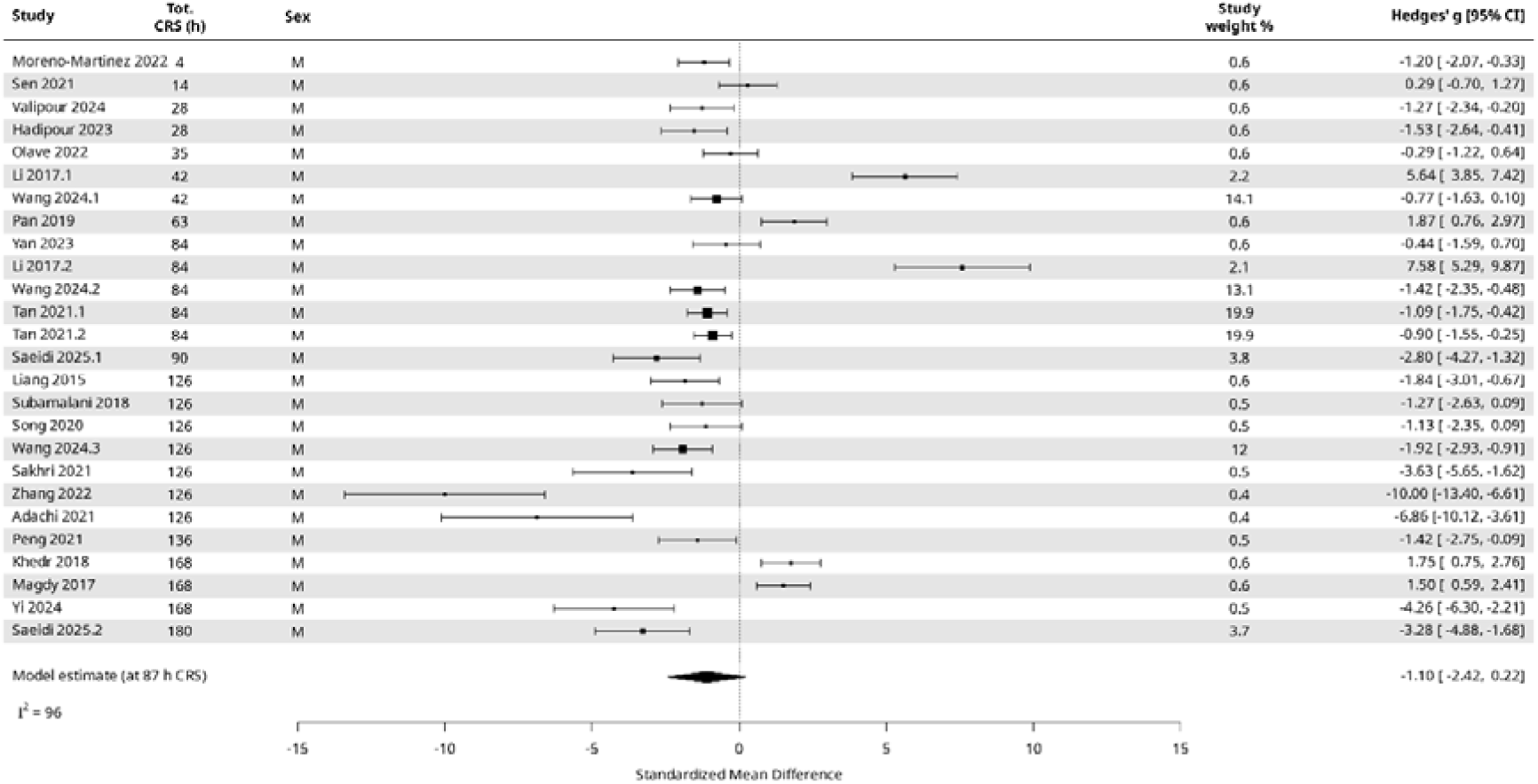
Forest plot of effect sizes extracted from studies measuring time spent in the central area of an OFT.

**Figure 8:**
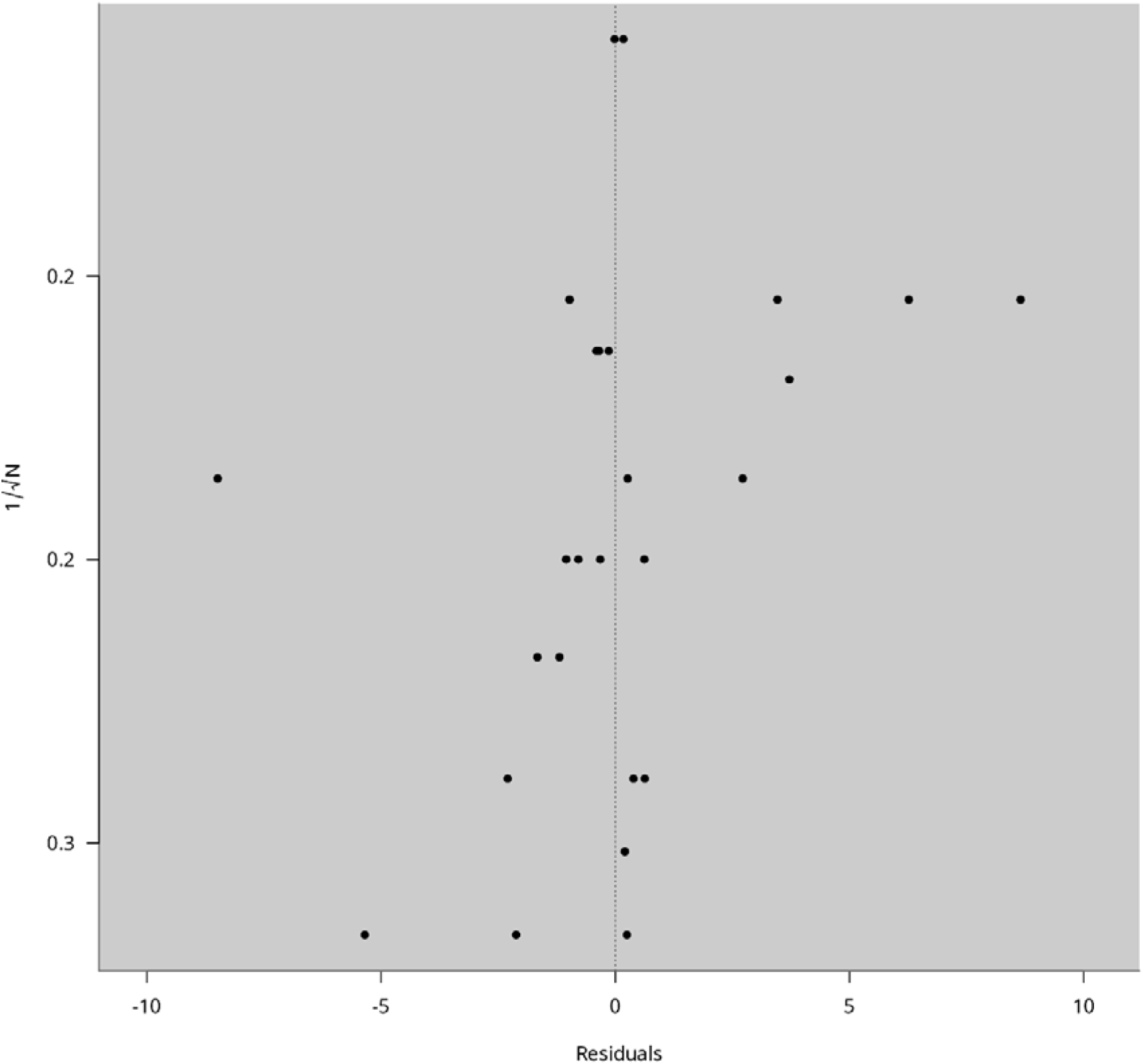
Funnel plot depicting effect size and study precision from studies measuring time spent in the central area of an OFT.

Across 51 studies (62 effect sizes), CRS was associated with a statistically significant decrease in the time spent in the open arms of the EPM (Hedges’ g =-1.41, 95% CI [-2.00,-0.83], t_60_=-4.828, p=9.87 x 10^-06^, calculated at the median CRS length of 63 hours). The total duration of CRS had a significant moderating effect on the time spent in the open arms (β_length_[95% CI] =-0.0097 [-0.0195, 0.0001], t_60_ =-1.981, p = 0.052) (Figure 9). Between-study heterogeneity was very high (I^2^ = 85.9%). Visual inspection of the funnel plot (Figure 10) showed evidence of asymmetry, indicating possible small-study effects.

**Figure 9:**
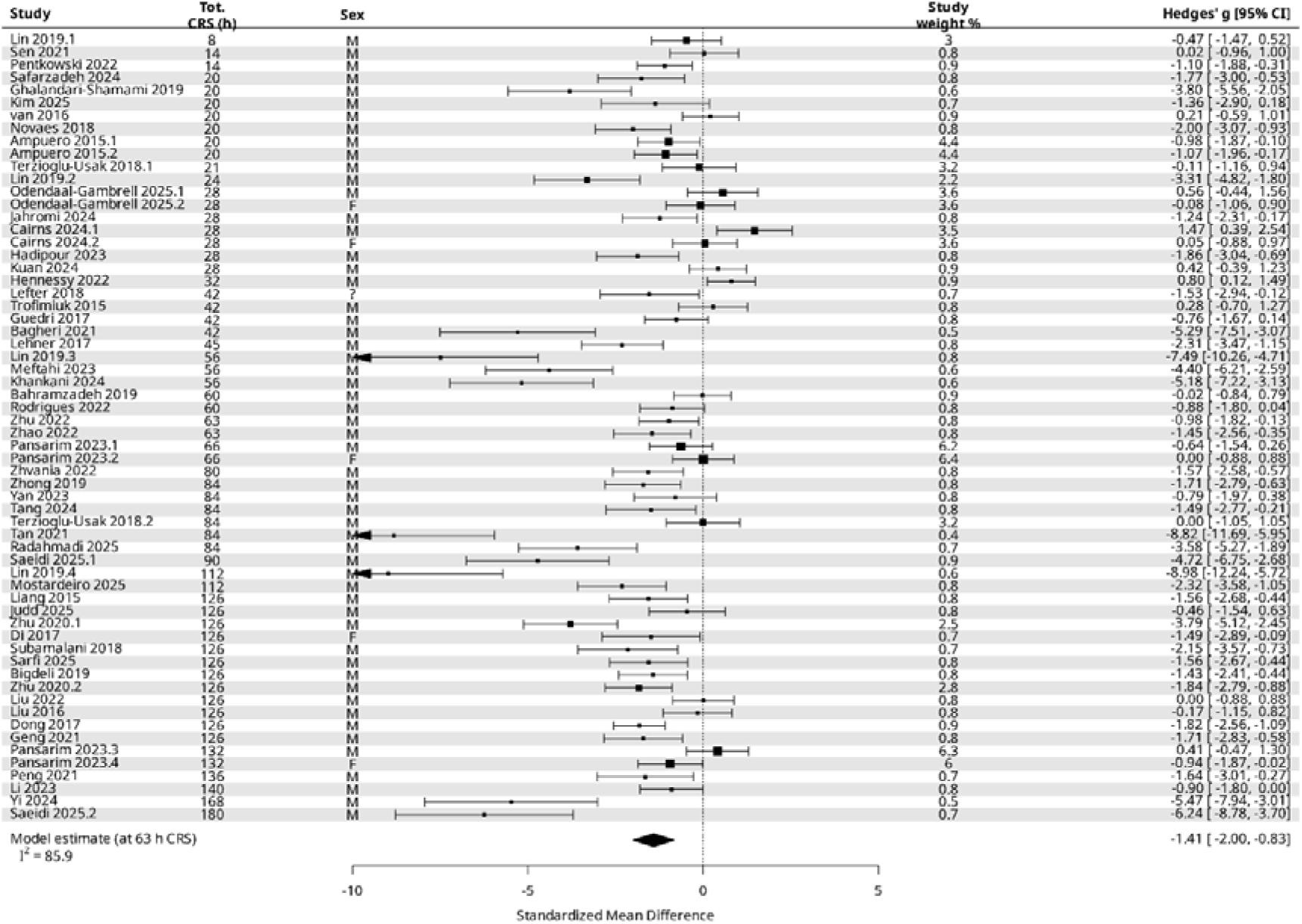
Forest plot of effect sizes extracted from studies measuring time spent in the open arms of an EPM.

**Figure 10:**
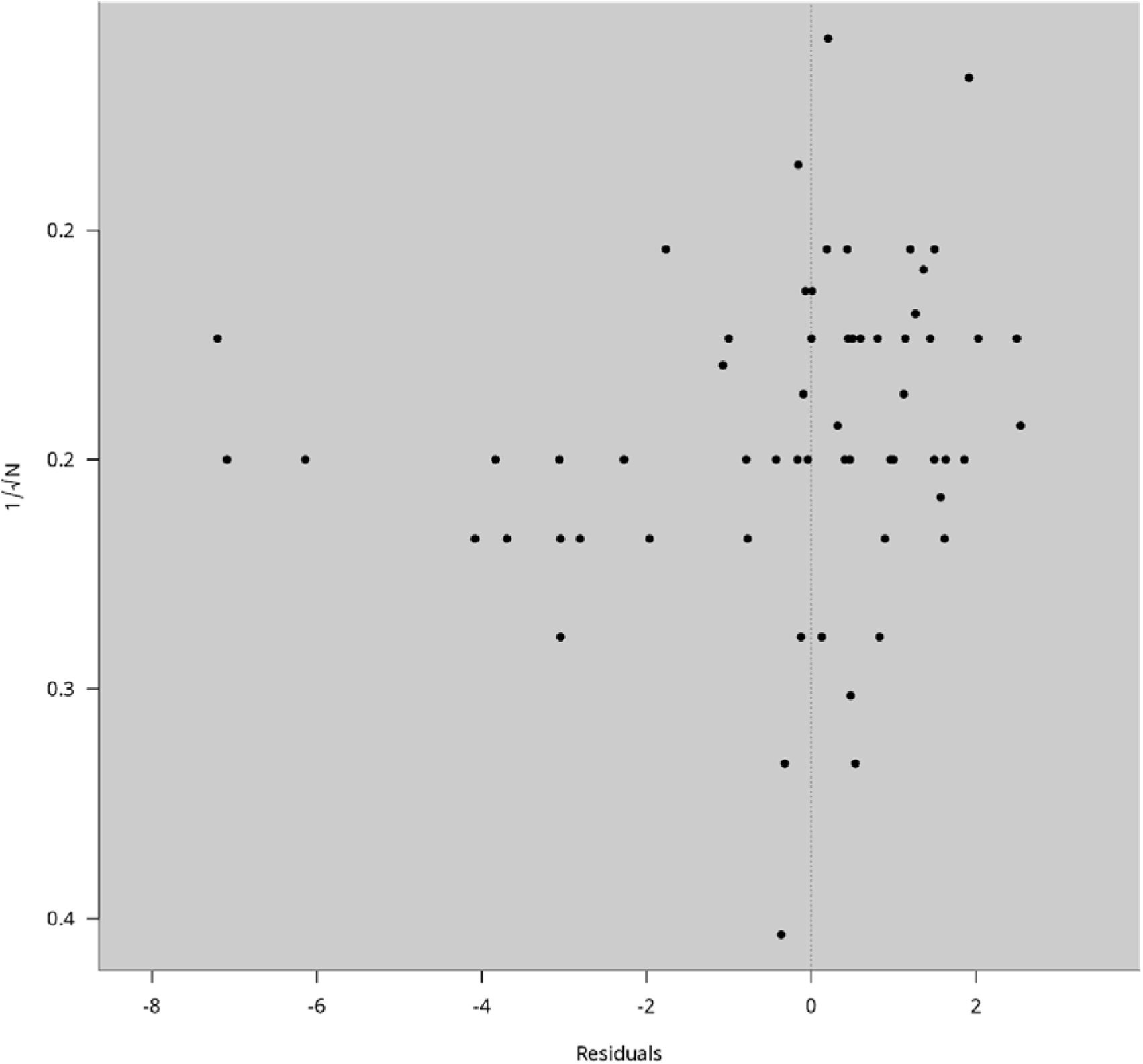
Funnel plot depicting effect size and study precision from studies measuring time spent in the open arms of an EPM.

In terms of correlations between tests, the strongest positive correlation was between the EPM and FST (Pearson’s correlation coefficient (r) = 0.78). There was also a positive correlation between the FST and SPT (r = 0.66), but SPT effect sizes correlated more weakly with those in the OFT and EPM (r = 0.39 and r = 0.51 respectively; Figure 11).

**Figure 11:**
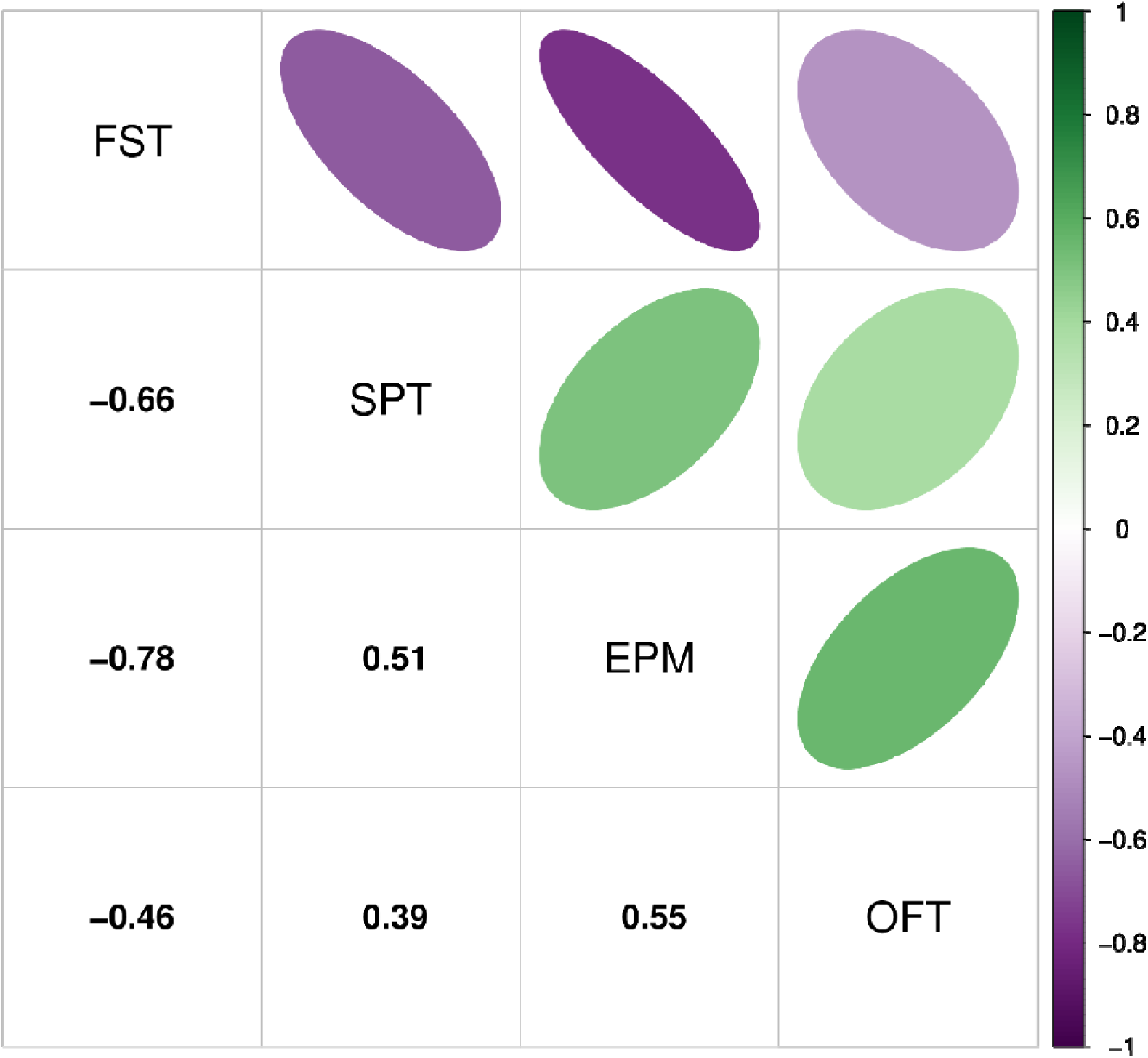
Directions and strengths of correlation between behavioural test effect sizes. Note that the negative values shown for FST correlations arise not because the magnitude of one effect decreases while the other increases, but because in one behavioural test the direction of the effect is positive and in the other it is negative.

A leave-one-out sensitivity analysis showed that no single article had a disproportionate effect on the predicted effect of CRS or on β_length_ or β_water_. These data are available at https://github.com/nicolaromano/Romano_Menzies_2026_CRS).

## Discussion

Our meta-analysis demonstrates that exposing rats to a CRS procedure increases immobility in the FST, decreases sucrose preference in the SPT, decreases the amount of time spent in the open arms of the EPM, but has no effect on the time spent in the centre of the OFT. Importantly, however, longer CRS procedures were not associated with larger effect sizes in any of these behavioural tests. Accordingly, our hypothesis that longer CRS procedures would be associated with larger behavioural effects was not supported. This suggests that (1) the intuition that a longer CRS procedure will cause a larger behavioural response across a panel of behavioural tests is incorrect, and/or (2) the behavioural methodologies studied here cannot reliably detect real differences in the behavioural consequences of CRS.

Our model generated effect size estimates (Hedges’ g) of 2.2 in the FST,-1.7 in the SPT, and-1.4 in the EPM. There is no clear consensus on what a biologically meaningful effect size is for these tests, and determining whether an effect size is biologically or clinically meaningful is not always straight-forward (Jordan *et al*., 2025). Cohen’s original classification of small (d⍰⍰=⍰⍰0.2), medium (d⍰⍰=⍰⍰0.5), and large (d ≥ 0.8) effect sizes is often cited, but is sometimes deemed problematic in the context of biology (Klein, 2005). We previously recommended selecting an effect size of d =-2.0 as biologically meaningful when determining a sample size for use in the SPT (Romanò & Menzies, 2025) because even “large” (d ≥ 0.8) effect sizes can be quite modest. For example, in one of the articles included in our present study (Wei *et al*., 2018), we estimated the SPT effect size to be-1.7, and this equated to a difference in sucrose preference of only 5% between the control and test groups after a three-week CRS procedure.

We note that our estimate of SPT effect size is lower than those of a recent meta-analysis that reported a standardized mean difference in post-CRS sucrose preference of-3.96 in Sprague-Dawley rats and-3.53 in Wistar rats (Mao *et al*., 2022). However, this difference may be explained by a systematic error in the way Mao et al. calculated some effect sizes (https://edin.ac/40XVwij).

Our effect size estimate of 2.2 for the FST in rats is in line with previous meta-analyses of FST immobility data in mice (Kara *et al*., 2018; Smalheiser *et al*., 2021), though in those studies the effect sizes related to the effect of anti-depressant treatments with respect to a baseline level of immobility, not our comparison between immobility in a stress-exposed group compared to a control group. A meta-analysis of behavioural tests carried out in rat models of neuropathic pain (which is presumably experienced by the animal as a chronic stressor) also reported effect sizes in the same range: Hedge’s g values of approximately 2.0 in the FST (and-1.2 in the SPT and-1.0 in the EPM; de la Rosa et al., 2024). However, we do not endorse use of the FST under any circumstances. This is because the FST’s face, construct, internal, external and predictive validity as a measure of stress-induced depression are in doubt (Nestler & Hyman, 2010; Molendijk & de Kloet, 2015; Anyan & Amir, 2018; Molendijk & de Kloet, 2019; Reardon, 2019; Carvalho *et al*., 2021; Trunnell & Carvalho, 2021; Ueno *et al*., 2022; Martins *et al*., 2025; Mohseni & Rafaiee, 2025), and the UK Government has recommended that the FST “*should not be used as a model of depression or to study depression-like behaviour…or for studies of anxiety disorders and their treatment*” (Anon, n.d.).

Our model predicted a modest effect size in the EPM (Hedge’s g =-1.4) that was not moderated by the total duration of restraint. We could not detect a significant effect of CRS on the time spent in the centre of the OFT (Hedges’ g =-1.1). This in line with other meta-analyses showing modest effects sizes in the EPM and OFT in stressed rodents (Wang *et al*., 2020; Çevik *et al*., 2025). Both the EPM and OFT measure whether an animal prefers to be in a protected or exposed area, with less time in the exposed area deemed to reflect a higher level of anxiety (Rosso *et al*., 2022), so it is perhaps unsurprising that effect sizes are similar. We found a strong positive correlation in effect sizes between these two tests in our previous study of behavioural tests after stressor exposure (Romanò & Menzies, 2025). However, we did not see such a strong correlation in the present study. We note that a large, multicentre initiative studying the replicability of the EPM found that data from only one of eleven original studies lay with the prediction interval derived from a meta-analysis of multiple replication studies, casting some doubt on the EPM’s usefulness (The Brazilian Reproducibility Initiative, 2025).

We noted a remarkable range of measurements were reported in articles using the OFT, but only a few of the included articles reported what we consider to be the conventional measure of anxiety: time spent in the centre of the OFT. A total of twenty-one different measurements were reported, ranging from the time spent moving and the distance moved to measurements of rearing, grooming, thigmotaxis and defecation. The design of the OFT in its current form(s) has been criticised (Stanford, 2007), citing issues in variability in the construction, size, shape and lighting of the arena, the duration of the test, whether and how long animals are adapted to the new environment, the animals’ previous experiences, their age and so on. Others have noted challenges associated with using the OFT (Walsh & Cummins, 1976; Paré & Glavin, 1986; Glavin *et al*., 1994; Gould *et al*., 2009; Prut & Belzung, 2003; Võikar & Stanford, 2023), including the possibility that the test in fact simulates a confrontation with a predator (the experimenter) alongside the simultaneous and unanticipated loss of social contact with cage-mates, and that the behaviour observed in the OFT represents “a compromise between responses aimed at minimizing detectability as a predator evasion tactic, and those which are motivated by attempts to reinstate contact with companions” (Suarez & Gallup, 1981).

We found that the strongest positive correlation between effect sizes was between the FST and EPM and between the FST and SPT; although the correlation between SPT and EPM effect sizes was only moderate. All other correlations were in or just below the moderate range, with the highest of these being between the EPM and OFT. Accordingly, our correlational analysis suggests that after CRS, the FST, EPM and SPT may all measure the same or similar consequence of CRS. In our previous meta-analysis of CVS articles (Romanò & Menzies, 2025), we also saw a very strong correlation between the FST and EPM (r = 0.91), but also between the FST and OFT (r = 0.94; compared to only 0.46 in the present study). In our CVS meta-analysis, the correlation between the FST and SPT was also fairly strong (r = 0.64), as was the correlation between the EPM and OFT (r = 0.74).

In articles that used more than one behavioural test (this comprises 58% of the total number of articles included in our study), we did not systematically document the order in which post-CRS behavioural tests were done, or the period of time between these tests. Accordingly, there may be order effects that we have not captured in our analysis. These order effects may be particularly pertinent if a behavioural test that is normally regarded as a stressor is performed prior to a subsequent behavioural test. For example, seventy-three articles included in our meta-analysis used the FST and at least one other behavioural test. A forced swim is sometimes used as a stressor in chronic variable stress protocols (Romanò & Menzies, 2025); it is plausible, therefore, that a behavioural test that takes place soon after the FST may quantify the animal’s on-going response to the FST as well as its underlying response to the CRS procedure.

Arguably more important than the absolute magnitude of our estimates of effect sizes was the finding that the effect sizes were not moderated by the total duration of restraint in any of the four behavioural tests. Before analysing the data, we speculated that observing a linear increase in effect sizes as CRS procedure duration increased may have been unlikely, and that we may observe an increase followed by a plateau or even an inverted-U-shaped curve if rats habituated to the stressor over time. Rats can adapt to repeated stressors: Buynitsky and Mostofsky (2009) point out that stress responses “will not necessarily continue to increase monotonically with the continued application of stressors. A peak may be reached and the animal may adapt to stress, causing [their stress response] to stop”. The authors suggested this may be due to an “insufficient intensity or duration of restraint”. Habituation and the associated blunting of hormonal and behavioural responses to homotypic restraint stress can emerge after a single week of CRS in rodents (Bhatnagar *et al*., 2002; Sadler & Bailey, 2016). This is consistent with the lack of a linear increase in effect size with length of stress in our analysis, though a blunting of effect size is not apparent from our analysis either. The molecular and cellular details of how the HPA axis decodes different stressors are complex and poorly understood; dishabituation can happen even during homotypic stress, for instance after a change in the duration of a single restraint episode (Kearns & Spencer, 2013). Extended periods of CRS may engage compensatory mechanisms that stabilize physiological activity to a different set-point, preventing further escalation of stress-related behaviours (Girotti *et al*., 2006; Herman, 2013). As a result, the measured behavioural effects may reflect early activation of stress responses and subsequent adaptation, rather than the cumulative duration of restraint exposure. We noted that the majority of the studies analysing the behavioural effects of CRS focused on behavioural measurements obtained at the end, or one or two days after the end, of the CRS procedure. We recently showed long-term transcriptional and functional changes in the HPA axis in mice exposed to CRS, that are visible only after the end of the chronic stress, and persist for several weeks in the recovery period (Duncan *et al*., 2024); while we did not perform behavioural testing in that study, these delayed molecular effects raise the question of whether comparable long-term adaptations might also occur at the behavioural level. An indication that this is the case comes from studies showing that a single 24-hour episode of restraint can induce long-term behavioural changes in mice that persist for up to five weeks (Chu *et al*., 2016).

The findings of our meta-analysis are reinforced by anecdotal evidence presented in a discussion of experiences with rats undergoing restraint in tubes as a refined (in the 3Rs sense) procedure for blood sampling: “rats were quite happy to crawl into the tubes, go to sleep and show no apparent signs that they had become stressed by the procedure”; “[Rats] will fall asleep given half the chance if being restrained for a few minutes or half an hour” (https://refinementdatabase.org/publications/enforced-restraint-of-rodents-a-discussion-by-the-refinement-enrichment-forum/). Similarly, one of us (NR) has observed mice voluntarily walking into restraint tubes at the beginning of a 30-minute period of restraint, and one study – carried out as part of a study exploring the motivation for rats to free a trapped cage-mate – provides evidence that rats may find having access to a restraint tube rewarding (Hachiga *et al*., 2020). However, another contributor to the discussion on refinements touched on the fact that these animals’ mental states are, obviously, not directly accessible to us, and although animals’ behaviours very likely do reflect their feelings, motivations and intentions, we may not always interpret these behaviours accurately: “It is my experience as clinical vet that animals are often very stoical, even though you know from clinical evidence that they are experiencing a lot of pain. This, actually, can make the work of a vet quite a challenge because the animal gives ‘wrong’ signals. Could it not be that a rat or a mouse who does not appear to be stressed while being immobilized in the tube, is in a state of anxiety and fear rather than sleeping?” (https://refinementdatabase.org/publications/enforced-restraint-of-rodents-a-discussion-by-the-refinement-enrichment-forum/).

Another possible factor that underlies different effect sizes after similar CRS procedures is *how* the post-CRS behavioural tests were carried out. In line with our previous observations (Romanò & Menzies, 2025), we documented variability in the design of given behavioural tests, and several other meta-analyses and narrative reviews of behavioural tests have critically discussed issues around variability in test protocols (Bogdanova *et al*., 2013; Kara *et al*., 2018; Brandwein *et al*., 2023; Verharen *et al*., 2023). For example, in the present study, we documented variability in the duration of the SPT (1-48 hours), the concentration of sucrose used (0.1-2%), and whether and for how long rats were food-and/or water-deprived before the start of the test (0-24 hours). We explored prior food/water deprivation as a potential moderator in the SPT but found no evidence for a systematic linear effect of deprivation. This is in contrast to another meta-analysis of the SPT that explored food and/or water deprivation prior to the SPT in rat models of chronic unpredictable stress that found longer periods of deprivation were associated with larger effects in the SPT, regardless of the duration of the stressor (Berrio *et al*., 2024). However, these authors argued *against* the use of food/water deprivation prior to an SPT because having food and water available is “the only scenario where one can be confident the observed changes relate to the hedonic value of the sweet, rather than to underlying metabolic drives”, and that “each hour of deprivation artificially inflates the observed effect, independently of stress, distorting the results and creating an overly optimistic picture of model’s success”. We agree with these sentiments, and because we could not detect an effect of food/water deprivation in our analysis, we echo their recommendation that deprivation is not used as part of an SPT procedure.

Turning to the other behavioural tests we studied; by visually inspecting the forest plots, we could not detect any obvious linear or non-linear effect of the total duration of restraint in the FST, EPM or OFT. We found that between-study heterogeneity was very high for all behavioural tests, indicating that there were marked differences in the animals used, the methods used and/or the way behavioural tests were carried out. If the total duration of restraint does not underlie differences in effect sizes in the FST, EPM or OFT, then which factors are influential?

The sex, age and/or strain of the animals may be important. For example, behaviour in the FST may be sex hormone-dependent in rats (Bogdanova *et al*., 2013; Kokras *et al*., 2015). However, the majority of articles included in our study used only males, and the age of rats used (when reported) was in a relatively narrow range of young adulthood. Almost all articles in our study used either Sprague-Dawley or Wistar rats and differences in their sensitivity to stress have been reported (Faraday, 2002; O’Mahony *et al*., 2011). However, we did not include strain as a moderator in our analysis.

We did not systematically extract data on how rats were housed in each of the included articles, but housing conditions may also affect rats’ behavioural responses to stress; some studies have shown that group-versus single-housing can impact the stress response in a sex-dependent way (Brown & Grunberg, 1995; Beck & Luine, 2002; Westenbroek *et al*., 2003). Given some researchers’ understandable concerns around these commonly used behavioural tests, an alternative approach is to investigate the effects of stressors by measuring behaviours that rodents spontaneously display in their home environment (Eraslan *et al*., 2023; Young *et al*., 2024). However, some of the more extreme types of behaviours observed under stress (for example, bar-biting, flipping and barbering) should not be thought of as ‘natural’ (Garner *et al*., 2004; Polanco *et al*., 2020; Ratuski *et al*., 2025). They are, by definition, abnormal and detrimental to the animal and its cage-mates (and, interestingly, can be mitigated by environmental enrichment; Gross et al., 2012).

Could the method of restraint be important? A range of restraint devices were used, and it is possible that rats perceive or respond to restraint differently, depending on the device used. Strict immobility (for example, complete immobility in a wire mesh) may be more stressful than restraint in a plastic tube, where the animal can readjust their body position, albeit to a limited degree. This question is not well-studied, and variability in the amount of detail provided in articles on how restraint is actually done makes this challenging to study systematically. However, there is some evidence that procedures that more strongly restrict the animal’s movement are more effective at inducing a stress response (Molina *et al*., 2023), and broader differences in how different experimenters handle, care for and interact with the animals during the study may also influence the animals’ behaviours in these tests.

Could the structure of the restraint procedure be important? The number of restraint sessions each day of the procedure was identical across most included articles (one restraint session per day), but the duration of individual restraint sessions ranged from one to eight hours per day. Because of this, the number and/or timing of restraint sessions per day may have influenced the impact of the stressor in a way that was independent of the total duration of restraint across the entire procedure (though Buynitsky and Mostofsky (2009) note that an “increase in the duration of restraint does not necessarily imply an increase in the stress experience”). In the articles included in our meta-analysis, the stressor was most commonly applied during the light phase (only four articles reported restraining rats during the dark phase). But rats are nocturnal and were, in many cases therefore, restrained at the time when they would normally already be less active or asleep. A further complication is that the activity of the HPA axis follows a marked circadian rhythm, with corticosterone levels being lowest during the light phase and peaking before the onset of the dark, active phase in rats. Consequently, the timing of restraint stress relative to this rhythm can influence both endocrine and behavioural outcomes. In rodents, this has been shown to be sex-and behaviour-dependent. For example, in mice, chronic stress applied during the dark phase increases anxiety behaviours selectively in females, while depressive behaviours are increased in males stressed during the light phase (Mingardi *et al*., 2025). Similarly, in male rats, chronic variable stress induces behavioural phenotypes related to anxiety and depression only when rats are stressed in the light phase (Aslani *et al*., 2014). The interpretation of these effects is further complicated by the fact that acute stress responses are stronger during the rising phase of the ultradian corticosterone secretion and smaller during the decaying phase (Windle *et al*., 1998; Sarabdjitsingh *et al*., 2010), and that chronic stress itself can change the basal level and rhythmicity of corticosterone secretion (Ottenweller *et al*., 1994).

To conclude, we cannot categorically recommend which test(s) should be used to evaluate the effects of CRS. However, one interpretation of our correlation analysis is that FST, EPM and SPT may all measure the same or similar consequence of CRS. Given the ethical issues around the FST and the relative simplicity of carrying out and quantifying the SPT, the SPT may be the best test to evaluate the effects of CRS. Specifically, a two-bottle choice paradigm over 12 hours during the dark phase of the light-dark cycle, ideally in group-housed rats, collecting data repeatedly over time in a within-subjects design. As before (Romanò & Menzies, 2025), we also call for the reporting of a fuller data set from SPTs: not just the % preference for sucrose, but also the amount of water (ml) and sucrose solution (ml and kJ) consumed, the amount (kJ) of food consumed, as well as regular measurements of the animals’ 24-hour food intake and body weight.

However, our meta-analysis provides no clear evidence on which to base a recommendation for an optimal duration of CRS. This is because the behavioural expression of a stress response may be more strongly influenced by other factors and, potentially, interactions between those factors. To investigate this further, it may be valuable to explore whether effect sizes in stress hormone measurements correlate with effect sizes in behavioural measurements. In this way, we can ask whether there is a relationship between the measurements normally used to define stress physiologically (i.e., activation of the hypothalamic-pituitary axis and increases in circulating cortisol/corticosterone and/or adrenocorticotropic hormone) and the behavioural tests that are believed to reflect stress-induced changes.

## Acknowledgements

This work was supported by the Medical Research Council [grant number MR/V012290/1]. For the purpose of open access, the authors have applied a Creative Commons Attribution (CC BY) licence to any Author Accepted Manuscript version arising from this submission.

